# Functional determinants of enhanced and depressed inter-areal information flow in NREM sleep between neuronal ensembles in rat cortex and hippocampus

**DOI:** 10.1101/264820

**Authors:** Umberto Olcese, Jeroen J. Bos, Martin Vinck, Cyriel M.A. Pennartz

## Abstract

Compared to wakefulness, neuronal activity during non-REM sleep is characterized by a decreased ability to integrate information, but also by the re-emergence of task-related information patterns. To investigate the mechanisms underlying these seemingly opposing phenomena, we measured directed information flow by computing transfer entropy between neuronal spiking activity in three cortical regions and the hippocampus of rats across brain states. State-dependent information flow resulted to be jointly determined by the anatomical distance between neurons and by their functional specialization. We distinguished two regimes, operating at short and long time scales, respectively. From wakefulness to non-REM sleep, transfer entropy at short time scales increased for inter-areal connections between neurons showing behavioral task correlates. Conversely, transfer entropy at long time scales became stronger between non-task modulated neurons and weaker between task-modulated neurons. These results may explain how, during non-REM sleep, a global inter-areal disconnection is compatible with highly specific task-related information transfer.

**Author Summary:** The brain remains active during deep sleep, yet we still do not know which rules govern information processing between neurons across wakefulness and sleep. Here we provide a first study of how information flow at the level of spiking activity varies as a function of brain state, temporal scale, brain area and behavioral task correlates of single neurons. We found that inter-areal communication at millisecond time scales is enhanced during sleep compared to wakefulness between neurons that code for task information. Conversely, non-modulated neurons showed more prominent communication at longer time scales. These results indicate that multiple, functionally determined communicative architectures coexist in the brain, and provide a novel framework to understand information processing and its consequences during sleep.

## Introduction

Wakefulness and sleep greatly differ in terms of neuronal activity and communication between brain areas (1–4). Since the pioneering studies of Moruzzi and Magoun (5), non-REM sleep (NREM) has been recognized as a complex and heterogeneously active brain state, which has been implicated in many functions ranging from memory consolidation (6) to synaptic homeostasis (7,8) and cellular restoration after intense activity (9). Crucially – and similarly to wakefulness – neural activity in NREM is not uniform throughout the brain: while slow wave activity is the hallmark of NREM in the neocortex (1), this type of sleep is also characterized by area-specific features such as hippocampal sharp-wave ripples (10) and thalamic spindles (11). Even within single anatomically defined areas, neuronal activity during sleep is not homogeneous, but is determined by factors such as the duration and the type of behavioral activity occurring during preceding wakefulness. For example, the intensity of slow wave activity in a given cortical area, as measured by EEG and local field potentials (LFPs), depends on the extent to which the area has been engaged in a task during wakefulness (12), to the point that individual cortical areas can enter slow-wave activity independently from the rest of the brain (13). A comparable dependence on behavioral activity was reported at the level of neuronal ensembles, as reactivation of spiking sequences occurs in the hippocampus, neocortex and subcortical structures during NREM sleep and quiet wakefulness (14–18). Task-dependent modulation of neuronal activity during NREM has been linked to sleep-dependent memory consolidation (12,19), although the underlying mechanisms are still a matter of debate. While reactivation of spike sequences during NREM (14,15,17,20) has been proposed as a way to consolidate previously acquired memories, slow wave activity has been hypothesized to mediate such consolidation via a global synaptic downscaling (21).

Surprisingly, limited attention has been paid to investigate how information flow between neurons is modulated across wakefulness and sleep, despite the fact that information transfer between neurons is often seen as a prerequisite for memory consolidation in both wakefulness and sleep (22–25). Most studies have been either limited to a mesoscopic level of analysis – and reported a drop in the ability of cortical areas to integrate information (3,26) – or focused on the coordinated reactivation of correlated spike trains both within the hippocampus and between the hippocampus and connected structures (14,17,27,28). However, a systematic attempt to characterize brain-state dependent modulation of neuron-level, directed information flow – both across spatio-temporal scales and between anatomically or functionally defined neuronal subpopulations – is lacking. Addressing this would solve one of the most puzzling questions concerning neural dynamics during NREM, i.e. how neuronal activity can be – at the same time – characterized by highly synchronous oscillatory dynamics [e.g. slow oscillations and sharp wave ripples (1,8,29–32)], and by the inability to properly integrate information across different areas. The most widely used approaches for computing pairwise relationships between neuronal activity – such as cross-correlograms (33) – are not able to discriminate between common sources modulating the activity of two neurons and actual causal links. Methods such as Granger causality analysis (34,35) do quantify directional links between time series, but only in terms of linear relationships, despite the fact that neural dynamics are highly non-linear (4,36). Here we addressed this issue by assessing transfer entropy (37) – TE, a non-linear counterpart of Granger causality (38) – between the spiking activity of pairs of neurons. This allowed us to quantify how directed information flow both within and between cortical areas [primary visual cortex (V1), barrel cortex (S1), perirhinal cortex (PRH) and hippocampus CA1 field (HPC)] varies between wakefulness and sleep at time scales ranging from a few up to hundreds of milliseconds. This was studied in rats trained to perform a sensory discrimination task in a maze, which allowed us to distinguish neurons that were either modulated by behavioral task elements or not. Our analyses reveal that: *a)* distinct communicative architectures operate at different time scales across brain states; *b)* relative to active wakefulness, inter-areal information transfer between hippocampus and cortex is enhanced during quiet wakefulness and NREM at the short (millisecond) time scale for neurons whose activity was modulated during the task; during NREM this increase is associated with sharp-wave ripples; *c)* conversely, inter-areal cortico-cortical information transfer is enhanced at long (hundreds of milliseconds) time scales during NREM between neurons which were not modulated by the task. These results challenge our current understanding of how neuronal communication varies across brain states by showing that long-range (i.e. inter-areal) neuronal communication can be not only preserved during NREM sleep, but also enhanced with respect to wakefulness, as a function of a combination of factors which cannot be considered independently: temporal scale, anatomical location and functional specialization during behavior.

## Results

### Spike-based Transfer Entropy

Transfer entropy (TE) was computed between spike trains of pairs of neurons using different temporal and amplitude binning, separately for the three brain states in which recordings were scored: active wakefulness (AW), quiet wakefulness (QW) and NREM [Methods, see also (4) and Fig 1A, B]. We computed TE for two subsets of temporal bin widths: 2-10 ms (compatible with interactions at the scale of a few synaptic steps), and 600-900 ms [in the range of low-frequency fluctuations in cortical activity (39), and in conformity with the range we previously employed to quantify non-linear correlations between such firing rate fluctuations across behavioral states (4)]. TE computed over short and long temporal bin widths will be called STE and LTE, respectively. TE has been shown to be a non-linear analogue of Granger causality (38), and as such is capable of quantifying both linear and non-linear directed information flow between two processes. To better understand the significance of STE and LTE in our dataset, we investigated the relationship between cross-correlograms and TE between neuronal pairs, computed for the three brain states under scrutiny (see Fig 2A-C for cross-correlograms and TE of an example neuronal pair). A significant correlation was observed between STE or LTE and the average value of the cross-correlogram computed, for the same neuronal pair, at the corresponding time scales (Fig 2D). Of relevance, the correlation becomes apparent when plotting both TE and cross-correlogram values on logarithmic axes, which is indicative of the non-linear relationship between TE and spiking patterns.

Besides being related to the cross-correlation between spike trains, TE is indicative of directed information flow (37). As such, it should capture asymmetries between values at the corresponding positive and negative time scales in the cross-correlogram. Such asymmetry has been attributed to spiking influences between neurons, both at short (29) and long (17) time scales. While at short time scales such influences correspond to direct synaptic connections (29,40), asymmetries at longer time scales are indicative of sequential (i.e. ordered) changes in firing rate (17,33). TE values should therefore allow to interpret correlations between spiking activity in terms of non-linear directional links. To assess the relationship between TE and cross-correlograms, we plotted the average STE/LTE value as a function of the cross-correlogram temporal bias index (33), for each directed neuronal pair *X*➔*Y*. The temporal bias index was calculated as *(P-N)/(P+N)*, where *P* and *N* represent the area under the cross-correlogram during, respectively, positive and negative time lags at the same temporal scales used for STE or LTE (Methods). Values of both STE and LTE were significantly different from 0 only for positive values of the temporal bias index (S1 Fig), corresponding to information flow along the direction accounted for by TE (i.e. from neuron *X* to neuron *Y*, see Materials and Methods). This remained true when focusing only on neuronal pairs showing significant STE or LTE values (Fig 2E), although we also found significant TE values for temporal bias indices between −0.5 and 0. This is indicative of the fact that TE is a non-linear estimator of information flow between spike trains. Non-linear effects play a minor role in determining the overall information flow when there are strong asymmetries in linear, rate-based information flow (i.e. for values of the temporal bias index <−0.5), but can become relevant for situations when there is only a mild asymmetry (e.g. for values of the temporal bias index between −0.5 and 0) – see Fig 2E and S1 Fig. In conclusion, directed information flow can be feasibly quantified in a non-linear manner by computing STE and LTE between neuronal pairs at multiple time scales.

**Fig 1.**
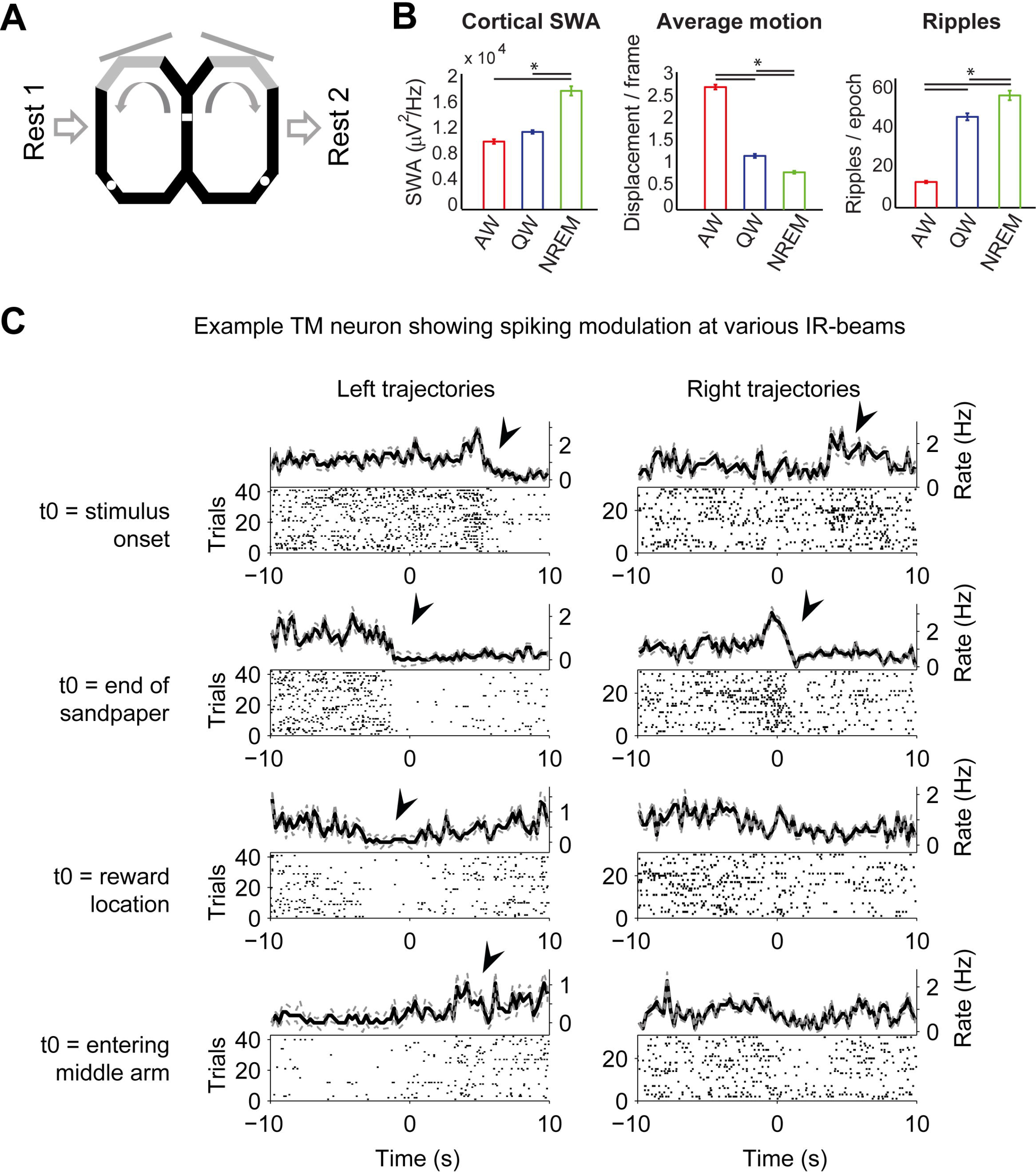
Analysis of brain states and behavioral task correlates. *(A)* Scheme of the experimental setup and design. Animals were trained to perform a visual discrimination task, flanked by two resting periods (Rest 1, Rest 2). White areas indicate reward locations. The white square indicates the location on the maze from where rats observed visual stimuli. Diagonal grey lines indicate the two screens. The grey areas on the maze show where the sandpaper was located. Details of the task can be found in Materials and Methods and in (4,62,63). *(B)* Scoring of brain states was based on visual inspection of LFP traces and body/head-motion patterns. A posteriori computation of slow wave activity *(SWA, left)*, average number of pixels of head displacement per frame *(center)* and occurrence of ripples *(right)* confirms that NREM was characterized by high SWA and low motion, and the opposite was true for AW. QW showed neither high SWA nor high motion. Error bars indicate SEM, significant differences were computed via one-way anova with post-hoc analysis. *(C)* Example PETHs for a TM neuron recorded in S1BF, for various infrared beams located on the maze, separately for trajectories on the left and right portion of the maze. Arrow heads indicate for which infrared beam crossings firing rate in the [−2 2] s period was significantly different than in the baseline period ([−10 −8] s range). Note how the PETHs show overlapping periods, for instance the decrease in firing rate in the top left PETH is the same that is then present in the PETH immediately below. This indicates that the whole task sequence was sampled by our analysis, and therefore that we could reliably detect neurons showing behavioral task correlates during recording sessions.

**Fig 2.**
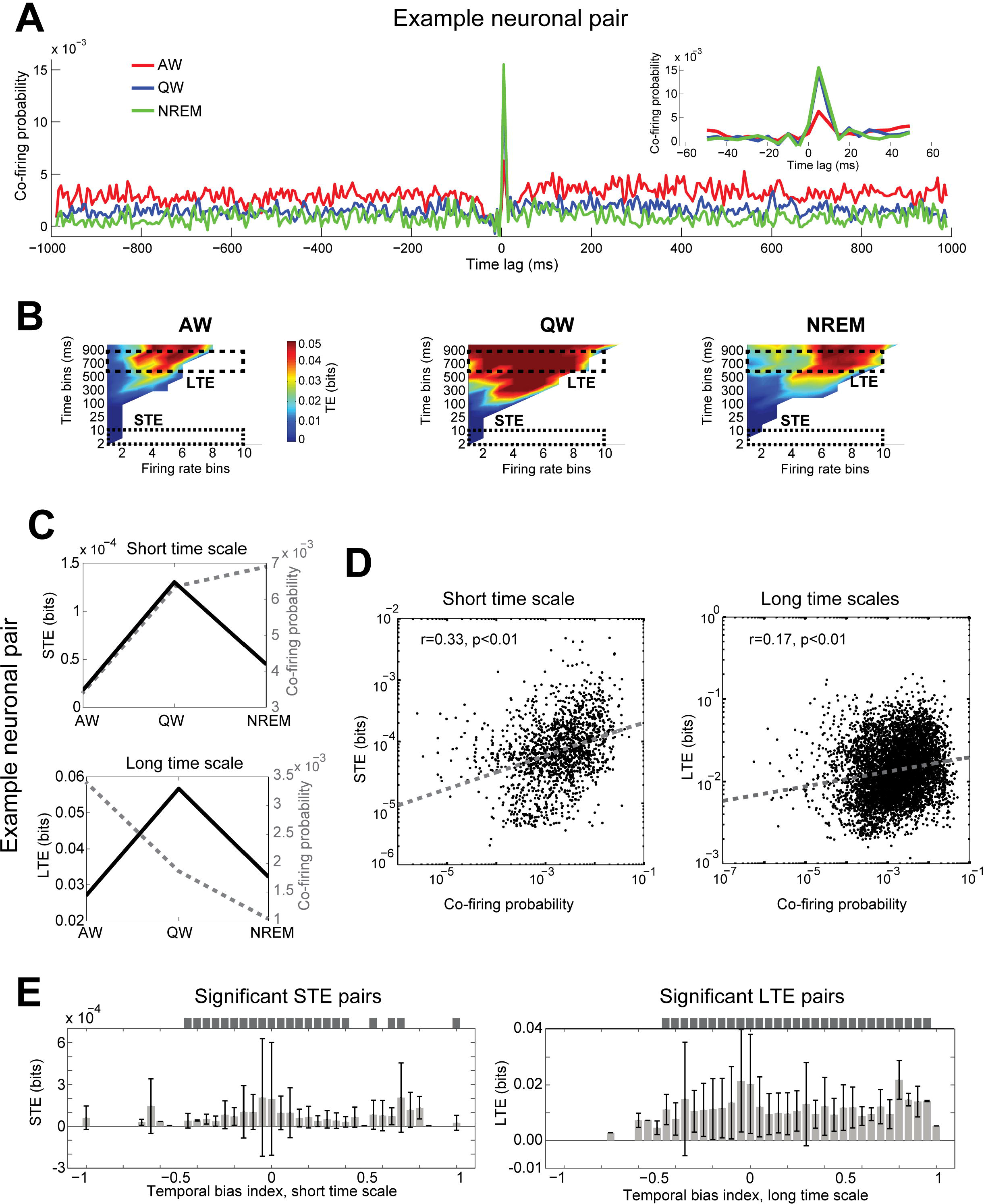
Interpretation of Transfer entropy (TE) as applied to spike trains. *(A)* Cross-correlogram computed for an example neuronal pair recorded in S1 (red: AW, blue: QW, green: NREM). Inset shows an enlargement of the [−60, 60] ms range. It can be seen that firing of the reference neuron in the pair is consistently followed by firing of the second neuron (at time lags in the [0, 10] ms range). This indicates an excitatory synaptic relationship. *(B)* TE computed during AW, QW and NREM for the neuronal pair shown in *(A)* for different time and amplitude bins (see Materials and Methods). Dashed rectangles indicate the ranges used to compute STE and LTE. White areas indicate ranges for which TE could not be calculated due to the lack of data points, and are thus not taken into account when computing STE and LTE. Note how values in the STE range, albeit lower than in the LTE range, are significantly higher than 0. *(C)* Comparison between TE (solid black lines) and co-firing probability (dashed gray lines) computed over short *(top)* or long *(bottom)* time scales, for the example pair shown in *(A, B). (D)* Relationship between TE values and average co-firing probability for short *(left)* and long *(right)* time scales. Each point represents one neuronal pair evaluated during one brain state, the three brain states are here superimposed; only points with positive TE values are shown. Negative TE values can occur as a consequence of the debiasing procedure (see Materials and Methods), and are always considered as null. Due to the non-linear relationship between TE and cross-correlogram of firing patterns, correlations between the two measures become especially evident when log-transforming the computed values (note the logarithmic axes). The dashed grey line in each plot indicates the result of the fitted linear regression. *(E)* Average STE *(left)* and LTE *(right)* values as a function of the cross-correlogram temporal bias index (Materials and Methods) of the corresponding neuronal pair, shown for pairs with significant STE/LTE values. Bins with TE values significantly higher than 0 are highlighted with a dark grey rectangle on top of the corresponding plot (independent samples t-test). Values in *(E)* are mean + SD.

### Short- vs. long-range information flow at different temporal scales

NREM has been related to a drop in functional connectivity with respect to wakefulness at time scales in the order of hundreds of milliseconds (2–4). In the TE measures, differences between wakefulness (either AW or QW) and NREM can be seen when considering average STE and LTE values for all recorded neuronal pairs (S3 Fig, left panels). For STE, there is a significant difference between AW and NREM (S3 Fig-A, left), while for LTE a difference is present between QW and NREM (S3 Fig-B, left). To better understand how spike-based directed information flow varies across brain states as a function of spatial and temporal scales, we computed STE and LTE separately for neuronal pairs located in the same or different brain regions.

Mean intra-areal STE showed – on average for all neuronal pairs – a marked decrease from AW to both QW and NREM, while no change was observed for mean inter-areal STE (Fig 3B, left). We further analyzed directed information flow only for neuronal pairs showing significant STE values, by computing the average STE value and the proportion of pairs with significant values for this subgroup. This analysis characterized whether the changes observed at the level of all neuronal pairs were due to variations in STE strength, to the number of neuronal pairs showing significant STE, or both. In terms of average STE values, significant intra-areal connections were stronger than inter-areal connections in all three behavioral states, and only intra-areal interactions presented a progressive decrease from AW to QW and NREM (Fig 3B, center). A different pattern was observed when quantifying the proportion of pairs showing a significant STE (Fig 3B, right). While no difference between the proportion of significant intra- and inter-areal connections was found during AW, the former became lower and the latter higher in both QW and NREM.

We next wondered if a similar pattern could be observed for LTE (Fig 3C-D). For intra-areal connections, we found lower LTE values in NREM than in both QW and AW, when considering all neuronal pairs (Fig 3D, left). This effect was the result of an increase in the strength of significant LTE values (which were higher in QW and NREM than in AW) and a progressive decrease in the proportion of significant LTE values (when going from AW to QW and NREM). A similar pattern was found for inter-areal LTE, except for a larger proportion of significant LTE connections in both QW and NREM than for intra-areal LTE. This resulted in an increase in average LTE over all neuronal pairs in QW compared to both AW and NREM, with no difference between AW and NREM (Fig 3D, left).

The results obtained so far do not show the expected, generalized drop in inter-areal connectivity reported during NREM both at the mesoscopic (3) and cellular scale (4). Instead, we only found a decrease for intra-areal TE, and a surprising increase in inter-areal STE. We next focused on investigating what could explain these results.

**Fig 3.**
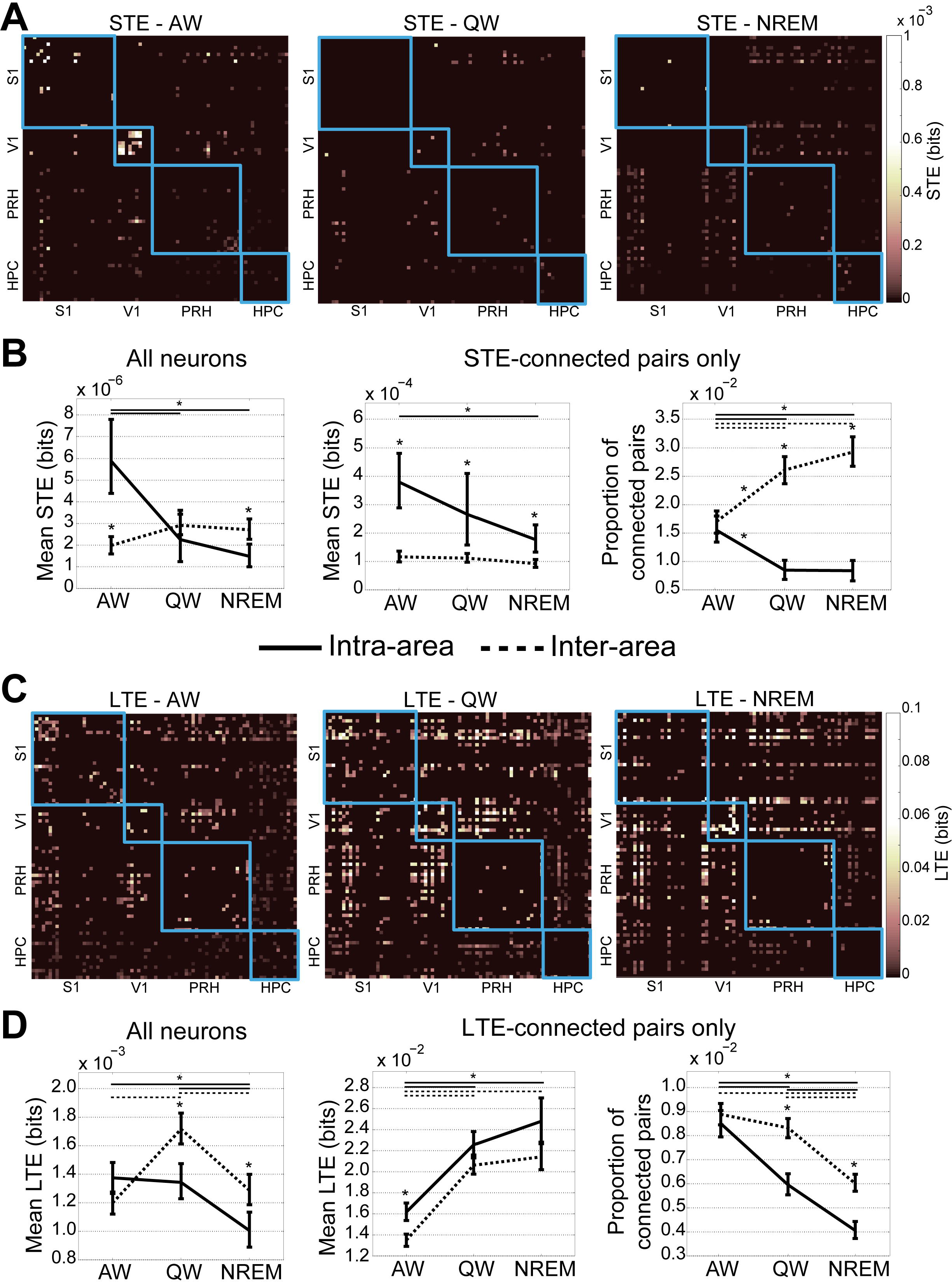
Brain-state dependent changes in intra- and inter-areal STE and LTE. *(A)* STE values for individual neuronal pairs computed for one example recording session across the three brain states. Values within blue boxes (also in the following Figs) refer to intra-areal connections. Points above the main diagonal indicate connections from neurons indicated on the vertical axis to neurons indicated on the horizontal axis, and vice versa for points below the main diagonal. *(B)* Average STE computed over all neuronal pairs *(left)*, or only pairs showing STE values significantly higher than 0 *(center)*, separately for intra-areal and inter-areal connections (solid and dashed lines, respectively). *(Right)* Proportion of neuronal pairs with significant STE. In this Fig and all following ones, error bars indicate bootstrap-estimated CIs (Materials and Methods). Asterisks on top of error bars indicate significant differences within a brain state, between intra- and inter-area connections. Asterisks on top of horizontal lines indicate significant differences between the brain states indicated by the line. The style of the line (solid or dashed) indicates to which group of neuronal interactions the asterisk refers. *(C)* Same as *(A)* for LTE. *(D)* Same as *(B)* for LTE.

### Role of behavioral task correlates in determining transfer entropy

In a recent study (4), we showed that neuronal coupling (measured by mutual information at time scales comparable with LTE) is not generally lower in NREM than in either AW or QW, but specifically between excitatory neurons located in different areas. For both STE and LTE, no major deviation from the observations reported for all neurons was found when we subdivided units into putative excitatory and inhibitory neurons (cf. Fig 2 and S4 Fig). We also analyzed whether the reported results could arise from differences between the various areas from which neurons were recorded, but none of the areas displayed major deviations from the general pattern (S5 Fig). Another factor that was previously shown to influence coupling between neurons was whether the spiking activity of individual cells was modulated by the task the animal performed during AW (4), here referred to as “task modulation”. We found that behavioral task correlates played a major role in determining STE and LTE during both QW and NREM. Based on this criterion (Fig 1C, Materials and Methods), neurons were subdivided into task-modulated (TM) and non-task-modulated (NTM) groups [Materials and Methods, (4)]. Strikingly, we found that most of the differences between intra- and inter-areal connections could be ascribed to the functional specialization of neurons.

For STE (Fig 4A-B), we found lower mean values in NREM than in AW for intra-areal connections both for pairs of TM and NTM neurons (Fig 4B, left). While no overall state-dependent change was observed for inter-areal STE, a significant difference appeared in NREM between TM and NTM neurons, with larger STE values for inter-areal connections between TM neurons. Strikingly, during NREM inter-areal STE between pairs of TM neurons was also stronger than intra-areal STE for pairs of either TM or NTM neurons. These state-dependent patterns were not due to variations in mean STE for significant STE pairs (Fig 4B, center), but rather to changes in the proportion of significant STE pairs (Fig 4B, right). Specifically, we found that, during QW and NREM, inter-areal STE between pairs of TM neurons showed a higher proportion of significant values than both inter-areal STE for NTM neurons and intra-areal STE for both TM and NTM neurons.

A different pattern was observed for LTE (Fig 4C-D). At the level of all neuronal pairs only behavioral task correlates (but not the intra- or inter-areal nature of connections) played a role in determining how LTE varies across behavioral states (Fig 4D, left). For LTE between pairs of TM neurons, we found a generalized drop going from AW to QW and NREM, which followed the decrease in the proportion of significant LTE values (Fig 4D, right), both at the intra- and inter-areal level. Conversely, mean LTE between all pairs of NTM neurons was higher in QW and NREM than AW, due to a combination of changes in the strength and proportion of significant LTE values. Moreover, the mean LTE strength for pairs of NTM neurons with significant LTE values was significantly higher for intra- than inter-areal pairs (Fig 4D, center). Overall, while during AW LTE was stronger between TM than between NTM neurons, the opposite was observed in NREM. For both STE and LTE, mixed connections (i.e. from a TM to a NTM neuron and vice versa) displayed values which were intermediate between those of TM and NTM pairs (not shown).

These results show that communication between neurons varies across behavioral states in a manner that is jointly determined by anatomical remoteness (intra- vs. inter-area), temporal scale and behavioral task correlates. Relative to AW, STE in NREM is characterized by an increase in the proportion of long-range information flow between TM neurons. In contrast, LTE presents a different, and almost opposite, pattern: during AW, there is a predominance in LTE between TM neurons, and the opposite is true for NREM, with stronger LTE between NTM rather than between TM neurons.

**Fig 4.**
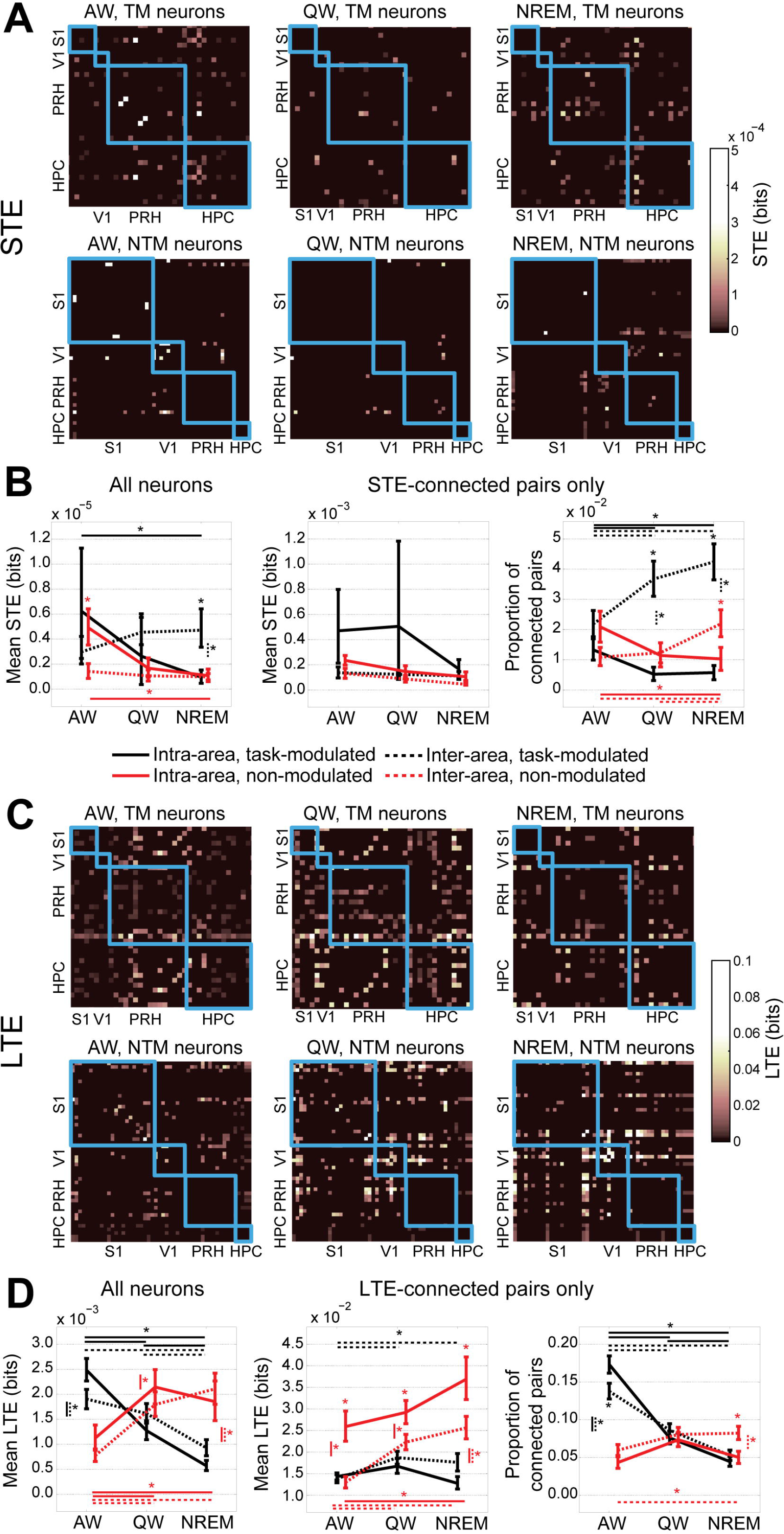
Role of behavioral task correlates in modulating STE and LTE across brain states. *(A)* STE for individual neuronal pairs computed for one example recording session across the three brain states. *(Top)* Connections between pairs of TM neurons. *(Bottom)* Connections between pairs of NTM neurons. *(B)* Average STE computed over all neuronal pairs *(left)*, or only pairs showing STE values significantly higher than 0 *(center)*, separately for intra-areal and inter-areal connections (solid and dashed lines, respectively) and for pairs of TM (black) and NTM (red) neurons. *(Right)* Proportion of neuronal pairs with significant STE. For within-state comparisons, asterisks on top of error bars refer to comparisons between pairs of TM and NTM neurons, for the same type of connection (intra- or inter-area). Asterisks next to a vertical line refer to comparisons between intra- and inter-area connections, for either TM or NTM neurons; the style of the vertical line indicates whether the comparison refers to intra- (solid) or inter-areal (dashed) connections, the color indicates whether values are higher for TM (black) or NTM (red) neurons. *(C)* Same as *(A)* for LTE. *(D)* Same as *(B)* for LTE. In all panels, plotting conventions are as in Fig 3.

### Potential mechanisms underlying the increase in inter-areal transfer entropy during NREM

The observed increase in inter-areal TE during NREM (between TM neurons for STE and between NTM neurons for LTE) is not in direct agreement with previous studies indicating a decrease in long-range communication during NREM with respect to wakefulness (3,4) – even if this was not always quantified by directional measures. To explore the determinants underlying this difference, we looked into differences between the individual brain regions from which neurons were recorded. We specifically focused on inter-areal connections during NREM, and investigated whether there were area-specific differences for pairs of TM and NTM neurons. As information transfer between HPC and cortical areas is thought to occur during NREM (6,41,42), we hypothesized that an increase in STE between TM neurons could be more prominent for connections between HPC and PRH or between HPC and primary cortices. Of relevance, we did not discern any significant directional difference between inter-areal STE and LTE values for any pair of areas (e.g. when comparing connections from HPC to PRH or vice versa – see also the Materials and Methods section), and we therefore always pooled the two directions together. Due to a smaller number of neurons recorded in S1 and V1 than in PRH and HPC – and no discernible difference between those two areas in the current analysis- we pooled S1 and V1 neurons together as cells recorded from primary sensory areas [PRIM, see also Materials and Methods, (4)].

We found that STE was stronger between TM neurons than between NTM neurons for connections between PRIM and HPC, or between PRH and HPC, but not for connections between PRIM and PRH (Fig 5A). This effect was due to a combination of changes in the strength and proportion of significant STE values (S6 Fig-A), and was highest for PRH-HPC neuronal pairs. For LTE, conversely, we found a strengthening of connections between PRIM and PRH for NTM neurons, but no change for connections involving one cortical area and HPC, for both TM and NTM neurons (Fig 5B, S6 Fig-B). Complete results for all brain states can be seen in S7 Fig.

Sharp-wave ripples (10) are an oscillatory phenomenon of hippocampal origin that has been implicated in mediating sleep-dependent consolidation of declarative memories (27,28), by transferring information from HPC to cortical areas. We thus wondered whether ripples could play a role in enhancing STE in NREM sleep between cortical areas and HPC. We computed STE during NREM between individual areas using datasets from which ripple periods were excluded [Materials and Methods, (4)]. Importantly, we applied debiasing procedures (see Materials and Methods) to avoid the risk that differences in TE values were due to datasets with distinct durations (see also the legend of S2 Fig). When considering all neuronal pairs, a significant difference was found between complete NREM and non-ripple NREM for STE between PRH and HPC (Fig 5C), with lower STE when ripples were excluded from the analysis. The proportion of significant STE values was decreased when considering connections between both PRH and HPC or PRIM and HPC (S6 Fig-C); no difference was observed for connections between cortical areas (Fig 5C, S6 Fig-C). For inter-areal LTE no difference was found when considering datasets with or without ripples (S8 Fig-A).

Finally, we tested to what extent sharp-wave ripples were associated with the overall increase in inter-areal STE during QW and NREM (Fig 3B). When removing ripple periods, we found that STE between TM neurons was reduced with respect to STE between NTM neurons, which in turn was not affected by removing ripples (Fig 5D). This effect was most prominent for the proportion of significant STE values (S6 Fig-D). Surprisingly, although ripples are also present in QW (Fig 1B), no effect of ripple exclusion on STE was found during QW (Fig 5D, S6 Fig-D). Values of LTE were also not affected by removing ripples (S8 Fig-B). Because sharp-wave ripples during sleep have been implicated in the consolidation of recently stored memories (25,27,28), we also tested whether we could find differences in inter-areal STE between the NREM periods before and after task performance (Fig 1A). However, no difference was found in this comparison (S9 Fig).

Overall, these analyses indicate that the increase in inter-areal STE during NREM – compared to AW – between pairs of TM neurons is limited to communication between cortical areas and HPC and is for a large part accompanied by sharp-wave ripples. However, while the increase in inter-areal STE was also present in QW and was limited to the same sets of areas (not shown), only during NREM it was linked to the presence of sharp-wave ripples. Conversely, the inter-areal increase in LTE between pairs on NTM that we found during NREM – compared to AW – was limited to connections between cortical areas and was not associated to ripple activity.

**Fig 5.**
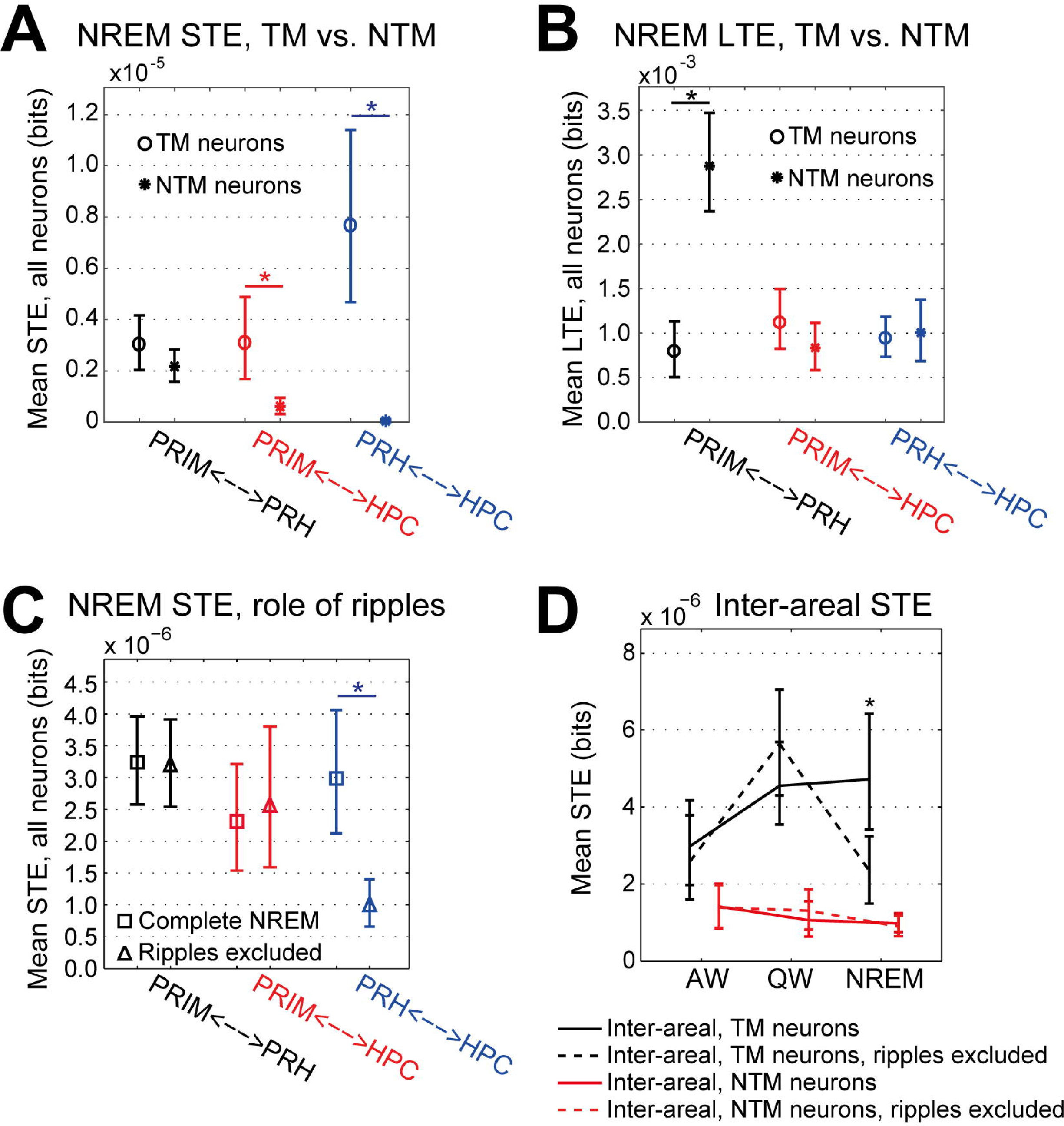
Regional dependence of changes in inter-areal TE during NREM. *(A)* Average value of inter-areal STE during NREM computed over all neuronal pairs, separately for connections between different brain areas (black: PRIM and PRH, red: PRIM and HPC, blue: PRH and HPC) and for pairs of TM or NTM neurons (circle and star shapes, respectively). Error bars indicate bootstrap-estimated CIs. *(B)* Same as *(A)* for LTE. *(C)* Same as *(A)* with STE computed over whole NREM datasets, or after removing ripple epochs (square and triangular shapes, respectively). (D) Average value of inter-areal STE across behavioral states, separately for pairs of TM or NTM neurons (black and red lines, respectively), and computed either on whole datasets (solid lines), or after removing ripple epochs (dashed lines). Error bars indicate bootstrap-estimated CIs. Note how removing ripples only affects STE between TM neurons during NREM. Asterisks indicate significant differences (p<0.05, bootstrap-estimated).

## Discussion

Understanding how neurons communicate during sleep, and how information flow is sculpted in different brain states is essential to uncover the mechanisms underlying sleep-dependent memory consolidation, but also to explain the loss of integrative functions characteristic of wakeful, conscious states (43,44). Generally, NREM has been associated with a decrease in the capability of neural systems to integrate information (3,4,45). An exception to this reduction in information transfer during NREM is represented by the sequential reactivation of stored memory traces across cortical areas and hippocampus (20,46–48), which has been hypothesized to contribute to sleep-dependent memory consolidation (6,14,15,41,49). This reactivation of memory traces is not a general property of all neurons within certain cortical structures, as it is selectively enhanced for recently acquired traces (14,15,17,20,46). How can these two apparently contrasting findings (decreased long-range communication and coordinated reactivation) be reconciled? Surprisingly, to our knowledge no study has attempted a consistent investigation of how communication between neurons varies across behavioral states and of which factors influence it. Here we provide evidence that behavioral task correlates are a major factor in determining long-range information flow across wakefulness and sleep, and can account for enhanced communication between specific neuronal subpopulations with respect to active wakefulness. Crucially, the influence of behavioral correlates is jointly determined by several factors: temporal scale of interaction, brain regions, underlying oscillatory dynamics and brain state. Specifically, STE between TM neurons in HPC and neocortical areas (and in particular PRH) is enhanced during QW and NREM with respect to AW. This enhancement is temporally linked to hippocampal sharp wave ripples during NREM but not QW. Conversely, LTE between NTM neurons is enhanced across neocortical areas (but not HPC) in NREM. Overall, our study shows how – during NREM – long-range communication can be both enhanced and depressed, based on time scale and the functional specialization of the involved neurons. At long time scales, compatible with cortical slow rhythms, inter-areal neocortical communication is specifically enhanced between NTM neurons, thus contributing to the maintenance of global neocortical coherence. During NREM TM neurons, on the other hand, engage in cortico-hippocampal communication at short time scales (Fig 6).

**Fig 6.**
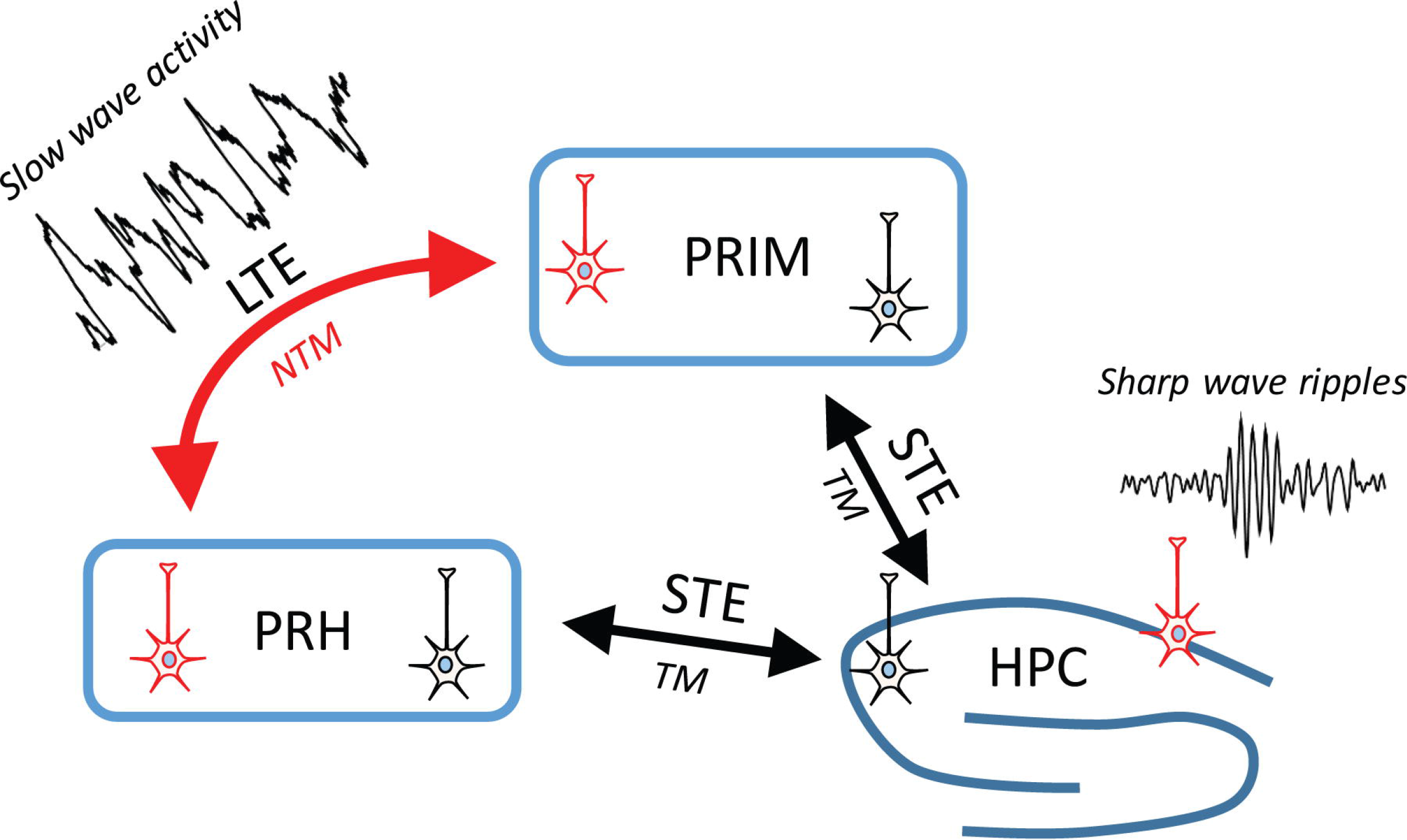
Schematic of inter-areal information flow during NREM. During NREM (as compared to wakefulness), STE is enhanced between TM neurons (black) located in HPC and neocortical areas; this increase in STE co-occurs with sharp-wave ripples. Conversely, LTE during NREM is enhanced between NTM neurons (red) located across neocortical areas, and follows the NREM-dependent increase in slow wave activity. Non-overlapping, functionally- and anatomically-defined neuronal subpopulations thus account for patterns of directed information flow at short and long time scales.

### Enhancement of inter-areal communication during NREM

During AW information transfer between neuronal pairs appears largely balanced, specifically across intra- and inter-areal connections and between TM and NTM neurons. However, during NREM we observed an increase in the proportion of significant inter-areal STE values between pairs of TM neurons relative to AW, and a general increase in LTE (strength and density) between NTM neurons (Fig 6). This cannot be fully grasped by only analyzing overall population-level dynamics, as the markedly different communicative architectures that we reported for neurons with distinct behavioral task correlates – and which co-exist at the same anatomical location – would average out. Furthermore, we showed the importance of taking into account temporal factors when assessing neural communicative architectures. Indeed, while STE reflects communication at the level of up to a few synaptic steps, LTE measures slower processing influencing the envelopes of activity – firing rate modulations (39) – that may be mediated by indirect, yet still causal, routes, such as spike-mediated traveling waves (30,50,51). These are changes in firing rate which operate over the course of tens or hundreds of milliseconds and rely on a coordinated modulation of spiking activity (39) rather than on temporally specific synaptic transmission between pre- and postsynaptic neurons. Importantly, the dynamics of slow oscillations during NREM [0.1 – 4 Hz (1,30,51,52)] – which are themselves traveling waves (30,53) – occur at time scales captured by LTE (600-900 ms).

### Potential mechanisms and functional role of the specific increase in inter-areal STE during QW and NREM

Recordings were performed in animals that had been trained daily over several months. It is therefore possible that interactions between TM neurons were strengthened with respect to those between NTM neurons. This would imply that the spontaneous activation of one TM neuron during QW or NREM would more easily propagate on a short time scale to other TM neurons. Surprisingly, however, this STE effect primarily occurs between rather than within brain areas (Fig 4B), and in particular between cortex and hippocampus (Fig 5, S6 Fig). This is in line with the hypothesis that – especially during NREM – memory traces are transferred and integrated across hippocampus and cortex (6), yet a mechanistic explanation for the increase in inter-areal STE is lacking.

First, it remains to be understood via which mechanisms the increase in STE is limited to inter-areal connections, despite the fact that cross-correlations and memory trace reactivation within HPC have been shown to increase in NREM following task performance (14,20,54). Previously, we showed that mutual information (a non-linear form of correlation) for firing rate modulations between individual neurons decreases for inter- but not intra-area pairs of excitatory neurons from AW to QW and NREM (24). This drop in long-range non-linear correlations might paradoxically underlie the increase in interarea (but not intra-area) STE, as during NREM weaker and slower synchronous changes in firing rate may allow the occurrence of spike-based interactions at shorter time scales. This might follow reduced noise correlations at longer time scales, which would otherwise override spiking interactions at short time scales.

Second, it needs to be addressed why the increase in inter-areal STE between TM neurons is limited to interactions between HPC and cortex. Sharp-wave ripples have been hypothesized as a potential mechanism mediating the consolidation of declarative memories via a coordinated transfer of information from HPC to cortex (10,27,54). Our data also indicate that high STE and ripples co-occur and are responsible for a major component of the increase in inter-areal STE during NREM (Fig 6). However, the co-occurrence of high STE and ripples is not present during QW, despite the fact that ripples can also be found in this state. Intriguingly, the role of ripples and memory trace reactivation in wakefulness seems to be different than in NREM, and is potentially not only linked to inter-areal information transfer mediating memory consolidation (55,56), but also to memory retrieval guiding decision making (57). It is thus possible that increased inter-areal STE between TM neurons during QW vs. NREM might be linked to distinct mechanisms and play different functional roles between the two brain states [e.g. memory consolidation in NREM, information retrieval or planning in QW (58–60)]. Strikingly, while ripple activity is present in both QW and NREM [albeit with different properties (10)], slow wave activity (SWA) characterizes cortical activity in NREM and might underlie the different mechanisms linking STE and ripples in QW and NREM. Indeed, coordinated, ripple-dependent memory trace reactivation between HPC and cortex has been reported to be maximal in the early sleep phase, during which SWA is strongest (14).

### Potential mechanisms and functional role of the increase in inter-areal LTE between NTM neurons in NREM

In contrast with STE, we observed an increase in inter-areal LTE between NTM neurons in QW and especially NREM with respect to AW (Fig 4C-D). Surprisingly, LTE between NTM neurons was generally stronger than that between TM neurons, and the increase in NREM was limited to connections between cortical areas. Combining these results with the STE data, it can be hypothesized that NTM neurons lack a task-induced strengthened coupling which can enhance STE and decouple neurons from underlying mass dynamics (see previous paragraph). Directed information flow between NTM neurons might instead be easily recruited by ongoing oscillatory activity capable of modulating firing rates at longer time scales (in the order of hundreds of milliseconds). Thus, while TM neurons can easily propagate precisely timed spike sequences (such as those elicited during AW or by fast population events such as ripples), directed information flow between NTM neurons is more readily deployed by slower population dynamics, hence their enhanced LTE during NREM in the cortex, in which SWA is at its highest.

How can this enhanced inter-areal LTE between NTM neurons be reconciled with the previously reported breakdown in long-range cortical connectivity during NREM (2–4,43)? Crucially, during NREM, cortical activity is at once disconnected yet highly synchronous, the latter due to the occurrence and propagation of slow oscillatory dynamics (1,30,51). Thus, decreased LTE between TM neurons (Fig 4D) might account for the reported drop in brain connectivity during NREM. Conversely, enhanced LTE between NTM neurons may provide a mechanism to maintain and propagate synchronous, slow firing rate fluctuations across the cortex (Fig 6). Future studies will be necessary to investigate this hypothesis, as the current literature on functional connectivity during brain states has not yet attempted to discriminate the neuronal sources of enhanced and disrupted coupling during NREM.

### Relationship with previous studies

Previous studies presented a somewhat contradictory picture of long-range neuronal dynamics during NREM, with both increased and decreased information flow between neurons. On the one hand, inter-areal communication was reported to be enhanced at short time scales – especially between the hippocampus and connected structures (14,17,20) – and is generally characterized by highly synchronous population activity, in particular in the neocortex (1,8). On the other hand, sleep and other brain states in which consciousness is lost have been characterized by a marked decline in the brain’s ability to integrate information between different areas (2–4,45). It must be pointed out, however, that most previous studies either did not focus on single neurons and did not explicitly discriminate between short- and long-scale temporal dynamics (3,61), or only assessed (undirected) correlations between the activity of single neurons (2,4). Conversely, STE and LTE quantify directed information flow between single neurons. By shedding light on how spiking information is transferred across brain states (millisecond- based spiking patterns for STE and long-scale fluctuations in spontaneous firing rates for LTE), our results thus greatly expand previous studies, for example by indicating circuit-level explanations for the different patterns of neuronal activity that are present in distinct brain states.

## Conclusions

Our study presents a novel picture of neuronal dynamics across brain states, which challenges traditional concepts of how communication between single neurons is modulated from wakefulness to sleep. By going beyond average population-level dynamics, and by incorporating in our analyses factors such as temporal scale, anatomical localization and behavioral task correlates, we were able to show how information flow differentially varies across behavioral states depending on the combined effect of all aforementioned factors. Specifically, we showed that neural correlates of behavioral task elements play a role comparable to anatomical factors in determining how neurons transfer information across brain states, and that distinct communicative architectures coexist at different time scales. Task-related and unrelated neurons thus obey different regimes for information transfer during wakefulness and sleep. Future studies will be necessary to understand the functional consequences of these results. For instance, it will be important to link the changes in STE and LTE for distinct functionally defined neuronal subpopulations to different forms of sleep-dependent memory processing, to expand and refine our study to other brain regions, neuronal subpopulations and brain states (e.g. REM sleep, extended recordings over the 24-hour wake-sleep cycle), and to study the functional consequences of enhanced STE/LTE by experimentally manipulating their values.

## Materials and Methods

The data utilized for this study was collected during experiments described in recently published manuscripts (4,62,63). For this reason, while all analyses presented here have been performed independently and do not overlap with the results presented in those articles, methods and data preprocessing will be only summarized here. All analyses were performed via custom-made scripts developed in Matlab (The Mathworks, Inc.).

### Subjects

>All animal experiments were conducted according to the National Guidelines on Animal Experiments and were approved by the Animal Experimentation Committee of the University of Amsterdam. Data was collected from three male Lister Hooded rats (28-46 weeks, obtained from Harlan), which were kept in a reversed day/night cycle (lights off time: 8:00 a.m., lights on time: 8:00 p.m.). Animals were food restricted so that their body weight was maintained at 85% of that of ad libitum fed animals (as per Harlan growth curves). Water was available ad libitum during all experimental phases.

### Behavioral setup

Rats were trained to perform a two-choice visual discrimination task on a Fig-eight maze (Fig 1A). Animals were trained to discriminate visual stimuli presented simultaneously on two monitors (grey diagonal lines, Fig 1A). Stimuli began after animals had crossed an infrared beam in front of the movable transparent door that was closest to the monitors (white square, Fig 1A). Animals were trained to choose the side arm of the maze (either left or right) corresponding to the screen where the positive conditioned stimulus (CS+) was shown (a specific white shape appearing over the black background); a CS- stimulus was shown on the other screen. After the door was lowered, if animals chose the correct side of the maze they were rewarded with two or three sugary pellets, which were placed in a ceramic cup (white circles in Fig 1A). Upon entering one side of the maze, animals would encounter strips of sandpaper on the walls of the maze (grey areas in Fig 1A). Visual stimuli ended when animals crossed an infrared beam located at the end of the area where sandpaper was present. The grain of the sandpaper predicted the amount of reward animals would receive after choosing the correct side arm (62,63). One pellet was also provided in the middle arm of the maze if animals had chosen the correct side arm in the previous trial. Eight infrared beams were employed to synchronize animal position to electrophysiological recordings, and to control the behavioral setup via custom-made software. Here we were only interested in discriminating whether the firing rate of neurons was significantly modulated along the various phases of the task. Thus, we did not discriminate between correct and incorrect trials. All animals were highly trained on the task and performed 60.1 ± 18.7 (mean ± standard deviation) trials per recording session, with an average success rate of 59.2 ± 0.9 % (mean ± standard error), significantly different from chance (p=2.1×10^−8^). For further details on the task, see (62,63).

Each recording session started and ended with a resting phase during which animals were placed in a flower pot on top of the maze (Fig 1A) under dim lights. Each resting phase usually lasted 0.5-1 hour. Task performance lasted approximately 1.5 hours, during which animals performed the behavioral task in dim light. The two resting phases were pooled together; no significant difference in STE or LTE was observed between them (p>0.05).

### Surgical procedure and recording drive

The right brain hemisphere of each rat was implanted with a custom-built tetrode microdrive, with 36 individually movable tetrodes (4,62,63). Eight recording tetrodes were directed to the monocular portion of the primary visual cortex (V1M, −6.0 mm posterior and −3.2 mm lateral to bregma), eight to hippocampal area CA1 (HPC, −3.5 mm posterior and −2.4 mm lateral), eight to the barrel field of the primary somatosensory cortex (S1BF, −3.1 mm posterior and −5.1 mm lateral) and eight to the perirhinal cortex (PRH, −5 mm posterior and −5 mm lateral, with an angle of 17 deg relative to the skull midline). One additional tetrode per area was used as a local reference. Tetrodes were slowly lowered from the brain surface towards the areas of interest, and then advanced daily to record different neurons at each recording sessions. Histological reconstructions were performed to localize tetrode positions after the final recording session. For details, refer to (62,63).

### Data acquisition and pre-processing

Data was acquired at 32 kHz via a Digital Lynx system (Neuralynx). Signals were filtered in the 1-500 Hz range to obtain local field potentials (LFPs). LFPs and motion tracking (64) were used to manually score behavioral states. To this aim, recording traces were subdivided into 4s-long epoch, and each was classified as AW, QW and NREM (4). All other behavioral states were excluded from the current analyses. Signals were filtered in the 600-6000 Hz range to perform spike detection. Action potentials were automatically extracted whenever the signal on any of the leads of a tetrode crossed a pre-set voltage threshold (4). Action potentials were assigned to single neurons by using a semi-automated spike sorting algorithm [see (24,62,63) for details]. Video acquisition of rat behavior was performed at 25Hz and was synchronized to electrophysiological recordings. Motion tracking of the headstage was performed as previously described (4). Interruptions of infra-red beams were monitored at 32 kHz.

Data were recorded during a total of six recording sessions in three different rats (24) (two recording sessions per animal). Each recording session was characterized by at least four neurons in each recorded region, and at least 30 minutes were spent in each state (AW, QW and NREM). On average, we recorded 13 ± 4 neurons from S1BF, 6 ± 2 neurons from V1M, 24 ± 10 neurons from PRH and 22 ± 4 neurons from HPC. Behavioral scoring resulted, on average, in 6293 ± 543 s of AW per session, 2277 ± 169 s of QW, and 1774 ± 390 s of NREM. All numbers are represented as average ± SEM.

### Behavioral state scoring

Scoring of behavioral states was performed manually, following standard methodologies applied to rodent behavior, based on cortical LFP and motor activity. The exact methodology has been previously described (4), and results in a subdivision of the recording sessions into 4-second-long epochs of either active wakefulness (AW), quiet wakefulness (QW), or NREM sleep (NREM) (see also Fig 1B). Any epoch that did not fulfil the criteria for AW, QW or NREM was discarded. Only sufficiently long periods of AW, QW or NREM (defined as periods lasting at least 30 s) were taken into account. As only a limited amount of REM sleep was present in our dataset, REM sleep epochs were not taken into account for further analysis.

### Determination of putative excitatory and inhibitory neurons

Neurons were subdivided into putative excitatory neurons and putative inhibitory neurons on the basis of their action potential waveforms (4,65). 66/381 Recorded neurons were classified as putative inhibitory fast-spiking neurons, corresponding to 17.3% of all recorded neurons and in agreement with previously recorded proportions of inhibitory neurons in cortex and hippocampus (29,65,66).

### Determination of task-modulated and non-modulated neurons

Peri-event time histograms (PETHs) were constructed for individual neurons based on the times each of the eight infrared beams present on the maze was crossed by animals during task performance (see Fig 1C for an example). Neurons were classified as task-modulated (TM) if their firing rates in the response window ([−2 2] s range around the crossing of a beam) were significantly higher or lower than the average firing rate in the baseline window ([−10 −8] s range around the crossing of a beam) (4). Out of 381 recorded neurons, 192 were classified as TM.

### Detection of hippocampal ripples

An automated procedure was implemented to detect ripples in the hippocampal LFP signals, based on typical spectral and temporal features of these oscillatory events (10,17,49). Details are reported in (4).

### Detection of up and down states during NREM

Up and down states were detected based on the approach presented in (67) and previously used in (4) (online-only supplemental material). The LFP signal from S1BF was filtered in the slow wave and delta range (0.5-4.0 Hz), and the instantaneous phase was computed via the Hilbert transform. A normalized multi-unit activity (MUA) measure was then computed combining all spike trains from neurons isolated from S1BF into a single, 1 ms bin, spike train. The resulting MUA trace was binarized (MUA:bin), to base the analysis of up and down states only on the presence or absence of action potentials. We then computed the circular histogram of MUA:bin firing rates as a function of the phase of the low-passed filtered LFP (4). Following (67), we computed the chance of an up state being observed as L(t)=cos(ϕ(t)- θ), where ϕ(t) is the instantaneous phase of the band-pass filtered LFP and θ is the phase corresponding to the peak in firing rate. The distribution of L values was subdivided into three segments (corresponding to up states, down states and undetermined transition states) by using *k*-means clustering. Only up and down states with a duration of at least 200 ms were considered in our analyses. More details about the detection of up and down states can be found in [(4) - online-only supplementary material].

### Measurement of directed information flow

Spike-based information flow between binned spike trains for pairs of neurons, quantified as a function of brain state, was measured using transfer entropy [TE (37,68)], according to the following formula:

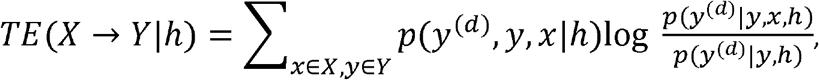

where *X* and *Y* are the sets of actual firing rates of two neurons (i.e. all firing rate bins); *x* and *y* are individual firing rate bins within *X* and *Y*, respectively; *h* indicates the behavioral state; *d* is the lag (in terms of time bins) between the activity of the neurons that are taken into account, and has been set to 1 in all analyses; *TE(X➔Y|h)* indicates the information flow from neuron *X* to neuron *Y* in a given brain state *h.* Firing rates were computed by binning spikes into non-overlapping bins. Bins were varied in terms of width (ranging from 2 to 1000 ms); firing rates were then subdivided into equipopulated bins (4,69); the number of firing rate bins was varied between 2 and 50 (4). The use of equipopulated binning ensured more robust estimates than the classically used method of equispaced binning (69) – although comparable values were obtained using this other binning method in preliminary analyses (not shown). TE values were debiased (to take into account the effects of finite sampling) via a shuffling procedure (4). Shuffled estimates were computed by scrambling the order of the time-binned firing rates of neuron *X* and also those of neuron *Y.* The same scrambling was used for *Y* and for its delayed version *Y(^d^)*, so that shuffling only affected estimates of causal information flow from *X* to *Y*, while influences of a neuron *Y* on itself were preserved. For each neuronal pair and brain state, we computed 10 shuffled STE or LTE estimates, and we subtracted their average value from that computed on the actual firing rates. This procedure was employed to reduce the bias in STE/LTE values due to limited sampling. Finally, we only considered the STE/LTE value to be significantly larger than 0 if the non-debiased estimate (i.e. the value obtained before substracting the shuffled estimate) also exceeded a significance threshold. This was computed as the bootstrap-estimated (100 repetition) 95^th^ percentile of the distribution of shuffled STE/LTE values. This last step enabled us to estimate the variability in STE/LTE values, independently for each neuronal pair, and thus only retain as significant those values exceeding such a noise threshold, thus increasing the robustness of our procedure to assess information flow.

Based on these computations, we focused on a subset of equipopulated amplitude bins (2-10 bins) and on two subsets of temporal bin widths (2-10 ms for STE and 600-900 ms for LTE). TE computed over short and long temporal bin widths will be called – respectively – STE and LTE. The values of temporal bins for LTE were chosen based on the range we previously employed to compute mutual information over long time scales (4). The values of firing rate (amplitude) bins were chosen empirically, in a range that resulted in non-zero values for a major portion of neuronal pairs, similarly to (4). For both STE and LTE, we computed the average TE value over the corresponding range of temporal and amplitude bins, and verified that this did not significantly change when ranges were varied by ±2 bins. We cannot exclude that different results could be obtained by increasing the lag term *d*, but an exhaustive analysis of this would be computationally challenging. Finally, we considered TE values to be significantly higher than 0 if they exceeded the 95^th^ percentile of the distribution of shuffled values (4). To further investigate – in addition to the analyses presented in Fig 2 – whether TE was related to the average firing rates of neurons *X* and *Y*, we employed a linear mixed-effects model to analyze the relationship between TE and average firing rates of neurons *X* and *Y* in different behavioral states. The correlation between LTE/STE and average firing rate of either neuron *X* or *Y* was not significant (p-value > 0.1). Conversely, the relationship between LTE/STE and the interaction term (computed via a linear regression) between the average firing rates of neurons *X* and *Y* was significant (p-value < 0.01). This confirmed that TE values are related to the interaction between the spiking activity of different neurons, rather than to their individual values.

We also verified how – during NREM – the computation of STE and LTE is affected by the presence of up and down states (1), as STE/LTE values could be specifically enhanced during up states. To this aim, we extracted NREM up and down states from NREM epochs [(67), see above for detailed methods], and computed STE/LTE on datasets from which we excluded – respectively – up or down states. We observed that excluding up states from the datasets invariably produced STE and LTE values not significantly different from 0 (S2 Fig). Excluding down states either did not significantly affect STE/LTE values or, for some pairs of brain areas, produced values not significantly different from 0 (which is likely due to a lack of sufficient data to reliably estimate TE). This control analysis indicates that TE values can be computed on whole NREM periods, as there is no underestimation of the information flow with respect to what could be obtained by excluding down (silent) states.

In spite of the precautions we took to properly estimate STE/LTE values, we did not detect significant directional differences between specific sets of areas. The lack of directional differences between TE computed between any pair of areas *(A, B)* is plausibly explained by the fact that the presence of recurrent (mono- or polysynaptic) connections can introduce causal influences in both directions (i.e. both from *A* to *B* and from *B* to *A)*. Also, we did not find an effect of task performance during AW on enhancing STE between TM neurons (S9 Fig) – as could be expected from previous literature (14,15,17,20,33,49).

### Cross-correlograms

Cross-correlograms were computed, for each directed neuronal pair *X*➔*y*, by calculating, for each action potential generated by the reference neuron *X*, the probability of a spike being generated by neuron *Y* in the [−1000, 1000] ms range, subdivided into 2 ms bins. The temporal bias index (33) was computed as *(P - N)/(P + N)*, where *P* and *N* are, respectively, the average value of the cross-correlogram (i.e. the firing probability of neuron *Y* following a spike by neuron *X)* in the positive (*P*) or negative (*N*) temporal range corresponding to that used to compute either STE or LTE, as appropriate (2-10 ms for STE and 600-900 ms for LTE).

### Statistical analyses

A bootstrapping procedure was used to estimate confidence intervals (α=0.05) for both STE/LTE values and the proportion of STE/LTE values significantly higher than 0. For a given group of N pairs of neurons, confidence intervals were obtained by sampling with replacement a new set of N pairs of neurons, and computing the STE/LTE value or the proportion of significant STE/LTE values for the resampled subset. Resampling was performed independently for neuronal pairs belonging to different recording sessions. The procedure was repeated 1000 times, and the 2.5th and 97.5th percentile of the distribution of values were used as confidence intervals for the measured proportion. In case of comparisons involving only two groups of neuronal pairs or two conditions, non-overlapping confidence intervals were used as a criterion to assess significant differences (p<0.05). In case of multiple comparisons, the α value for assessing significant differences was adjusted using Bonferroni correction. In all Figs, displayed confidence intervals correspond to values computed using α=0.05.

## Acknowledgments

This work was supported by the European Union (EU FP7 ICT Grant 270108 “Goal-Leaders” to C.M.A.P.), the Netherlands Organization for Scientific Research (NWO ALW-Open Grant 820.02.020 to C.M.A.P.), by the EU Horizon 2020 program under Grant Agreement 720270, Human Brain Project SGA1 to C.M.A.P., by the FLAG-ERA JTC 2015 project CANON (co-financed by NWO) to U.O., and by the High Performance Computing and Networking Fund of the University of Amsterdam (project “GPU-based accelerators for SILS-CSN cluster”) to U.O. We thank Kenneth Harris (University College London; KlustaKwik) and A. David Redish (University of Minnesota, Minneapolis; MClust) for providing software, Jadin Jackson for help with animal training and recordings, Jan Lankelma for providing analytical tools, and the Technology Center of the University of Amsterdam.

## Supporting Information

**S1 Fig. Interpretation of STE/LTE values.** Average STE *(left)* and LTE *(right)* values as a function of the cross-correlogram temporal bias index (Materials and Methods) of the corresponding neuronal pair, shown for all neuronal pairs. Bins with TE values significantly higher than 0 are highlighted with a dark grey rectangle on top of the corresponding plot (independent samples t-test). Values in *(D)* are mean + SD.

**S2 Fig. Influence of up and down states on STE and LTE values.** *(A)* Average intra-areal STE during NREM computed over all neuronal pairs, separately for connections between different brain areas (black: PRIM, red: PRH, blue: HPC), and computed either on whole NREM datasets (circles) or after removing Up states (stars) or Down states (diamonds). Error bars indicate bootstrap-estimated CIs. *(B)* Same as *(A)* for inter-areal STE (black: PRIM and PRH, red: PRIM and HPC, blue: PRH and HPC). *(C,D)* Same as *(A,B)* for LTE. Overall, these results indicate that TE values computed over complete NREM epochs are not distinguishable from those computed over up states only, and show that no significant TE value could be computed for down states only. In some instances, null values were also found when computing TE on up states only (e.g. intra-areal STE in HPC), which can be ascribed to a reduction in the number of data points beyond that which is required to compute valid TE values.

**S3 Fig. Average STE and LTE computed over all neuronal pairs show a global net decrease in NREM compared to wakefulness.** *(A)* Average STE computed over all neuronal pairs *(left)*, or only pairs showing STE values significantly higher than 0 *(center). (Right)* Proportion of neuronal pairs with significant STE. Error bars indicate bootstrap-estimated CIs (Materials and Methods). Asterisks on top of error bars indicate significant differences between brain states. *(B)* Same as *(A)* for LTE.

The global net decrease in TE during NREM compared to wakefulness (between AW and NREM for STE, and between QW and NREM for LTE) can be seen when considering all recorded neuronal pairs (*left panels).* However, simply focusing on neuronal pairs with significant TE shows a largely different picture *(center and right panels).*

**S4 Fig. Neuronal subtypes do not explain differences between intra- and inter-areal STE/LTE connections.** *(A, Top)* Average STE computed for all pairs of putative excitatory neurons *(left)*, or only pairs of putative excitatory neurons showing STE values significantly higher than 0 *(center)*, separately for intra-areal and inter-areal connections (black solid and dashed lines, respectively). *(Right)* Proportion of pairs of putative excitatory neurons showing significant STE values. Error bars indicate bootstrap- estimated CIs. Significant differences are indicated following the plotting conventions of fig 4. *(A, Bottom)* Same as *(A, Top)* for LTE. *(B)* Same as *(A)* for pairs of putative inhibitory neurons (red lines).

**S5 Fig. Anatomical determinants of changes in STE and LTE across brain states do not explain the differences observed between intra- and inter-areal connections.** *(A)* Average STE computed over all neuronal pairs *(left)*, or only pairs showing STE values significantly higher than 0 *(center)*, separately for intra-areal *(top, solid lines)* and inter-areal *(bottom*, dashed lines) connections, and for individual pairs of areas (for color codes refer to S2 Fig). *(B)* Same as *(A)* for LTE. Error bars indicate bootstrap-estimated CIs (Materials and Methods). Asterisks on top of error bar indicate if (and which) set of areas (indicated by color) shows significantly higher or lower values than all other sets of areas in the panel, within a single brain state. Asterisks on top of lines connecting points for different brain states indicate significant differences between adjacent brain states. Asterisks next to a small horizontal line indicate significant differences between AW and NREM for a specific type of area-to-area connection (indicated by line color and style).

**S6 Fig. Anatomical and task-related determinants of the increase in inter-areal TE during NREM – supplementary data to Fig 5.** *(A)* Same as Fig 5A for the average STE computed during NREM only for neuronal pairs showing STE values significantly higher than 0 *(left)*, and for the proportion of significant STE values *(right)*, computed separately for TM and NTM neurons. *(B)* Same as *(A)* for LTE. *(C)* Same as Fig 5C for the average STE computed only for neuronal pairs showing STE values significantly higher than 0 *(left)*, and for the proportion of significant STE values *(right)*, computed separately for whole NREM datasets and after excluding ripple epochs. *(D)* Same as Fig 5D for the average STE computed only for neuronal pairs showing STE values significantly higher than 0 *(left)*, and for the proportion of significant STE values *(right).*

S7 Fig. Area-specific variations in TE patterns across brain states for TM and NTM neurons – supplementary data to Fig 5. *(A, top)* Average STE computed over all neuronal pairs *(left)*, only for pairs showing STE values significantly higher than 0 *(center)*, and proportion of significant STE values *(right)* for intra-areal connections (same color scheme as in S5 Fig), computed separately for pairs of TM and NTM neurons (solid and dashed lines, respectively). Error bars indicate bootstrap-estimated CIs. *(A, bottom)* Same as *(A, top)* for LTE. *(B)* Same as *(A)* for inter-areal connections. The same plotting conventions as in S5 Fig have been used. This Fig extends the data presented in Fig 5 to AW and QW. No statistical comparisons have been performed, as this Fig is only presented for exhaustivity. Statistical analyses for data related to NREM is presented in other Figs (Fig 5, S4 Fig).

**S8 Fig. LTE is not affected by ripple activity.** *(A)* Same as Fig 5C *(left)* and S6 Fig-C *(center, right)* for LTE. The same plotting conventions are used as in Fig 5C. (B) Same as Fig 5D *(left)* and S6 Fig-D *(center, right)* for LTE. The same plotting conventions are used as in Fig 5D. Confirming the finding that ripples are not related to LTE, no difference is visible between values computed either including or excluding them.

**S9 Fig. Effect of task execution on STE during NREM.** Average value of STE computed over all neuronal pairs *(left)*, only for pairs showing STE values significantly higher than 0 *(center)*, and proportion of significant STE values *(right)* for connections between PRH and HPC during NREM, computed during either Rest 1 or Rest 2. Error bars indicate bootstrap-estimated CIs. The presence of TE values significantly higher than 0 indicates that sufficient data is present to compute STE/LTE for Rest 1 and Rest 2 separately (cf. with S2 Fig).

## References

1. Steriade M, Timofeev I, Grenier F. Natural waking and sleep states: a view from inside neocortical neurons. J Neurophysiol. 2001 May;85(5):1969–85.

2. Peyrache A, Dehghani N, Eskandar EN, Madsen JR, Anderson WS, Donoghue JA, et al. Spatiotemporal dynamics of neocortical excitation and inhibition during human sleep. Proc Natl Acad Sci U S A. 2012 Jan 31;109(5):1731–6.

3. Massimini M, Ferrarelli F, Huber R, Esser SK, Singh H, Tononi G. Breakdown of cortical effective connectivity during sleep. Science. 2005 Sep 30;309(5744):2228–32.

4. Olcese U, Bos JJ, Vinck M, Lankelma JV, van Mourik-Donga LB, Schlumm F, et al. Spike-Based Functional Connectivity in Cerebral Cortex and Hippocampus: Loss of Global Connectivity Is Coupled to Preservation of Local Connectivity During Non-REM Sleep. J Neurosci. 2016 Jul 20;36(29):7676–92.

5. Moruzzi G, Magoun HW. Brain stem reticular formation and activation of the EEG. Electroencephalogr Clin Neurophysiol. 1949 Nov;1(4):455–73.

6. Diekelmann S, Born J. The memory function of sleep. Nat Rev Neurosci. 2010 Feb;11(2):114–26.

7. de Vivo L, Bellesi M, Marshall W, Bushong EA, Ellisman MH, Tononi G, et al. Ultrastructural evidence for synaptic scaling across the wake/sleep cycle. Science. 2017 03;355(6324):507–10.

8. Vyazovskiy VV, Olcese U, Lazimy YM, Faraguna U, Esser SK, Williams JC, et al. Cortical firing and sleep homeostasis. Neuron. 2009 Sep 24;63(6):865–78.

9. Vyazovskiy VV, Harris KD. Sleep and the single neuron: the role of global slow oscillations in individual cell rest. Nat Rev Neurosci. 2013 Jun;14(6):443–51.

10. Buzsáki G. Hippocampal sharp wave-ripple: A cognitive biomarker for episodic memory and planning. Hippocampus. 2015 Oct;25(10):1073–188.

11. Contreras D, Destexhe A, Sejnowski TJ, Steriade M. Spatiotemporal patterns of spindle oscillations in cortex and thalamus. J Neurosci. 1997 Feb 1;17(3):1179–96.

12. Huber R, Ghilardi MF, Massimini M, Tononi G. Local sleep and learning. Nature. 2004 Jul 1;430(6995):78–81.

13. Vyazovskiy VV, Olcese U, Hanlon EC, Nir Y, Cirelli C, Tononi G. Local sleep in awake rats. Nature. 2011 Apr 28;472(7344):443–7.

14. Ji D, Wilson MA. Coordinated memory replay in the visual cortex and hippocampus during sleep. Nat Neurosci. 2007 Jan;10(1):100–7.

15. Euston DR, Tatsuno M, McNaughton BL. Fast-forward playback of recent memory sequences in prefrontal cortex during sleep. Science. 2007 Nov 16;318(5853):1147–50.

16. Foster DJ, Wilson MA. Reverse replay of behavioural sequences in hippocampal place cells during the awake state. Nature. 2006 Mar 30;440(7084):680–3.

17. Lansink CS, Goltstein PM, Lankelma JV, McNaughton BL, Pennartz CMA. Hippocampus leads ventral striatum in replay of place-reward information. PLoS Biol. 2009 Aug;7(8):e1000173.

18. Pezzulo G, van der Meer MAA, Lansink CS, Pennartz CMA. Internally generated sequences in learning and executing goal-directed behavior. Trends Cogn Sci. 2014 Dec;18(12):647–57.

19. Rasch B, Büchel C, Gais S, Born J. Odor cues during slow-wave sleep prompt declarative memory consolidation. Science. 2007 Mar 9;315(5817):1426–9.

20. Wilson MA, McNaughton BL. Reactivation of hippocampal ensemble memories during sleep. Science. 1994 Jul 29;265(5172):676–9.

21. Tononi G, Cirelli C. Sleep function and synaptic homeostasis. Sleep Med Rev. 2006 Feb;10(1):49–62.

22. Hebb DO. The Organization of Behavior: A Neuropsychological Theory. [Internet]. Psychology Press; 1949 [cited 2017 Mar 29]. Available from: https://www.amazon.com/Organization-Behavior-Neuropsychological-Theory/dp/0805843000

23. Dan Y, Poo M-M. Spike timing-dependent plasticity of neural circuits. Neuron. 2004 Sep 30;44(1):23–30.

24. Olcese U, Esser SK, Tononi G. Sleep and synaptic renormalization: a computational study. J Neurophysiol. 2010 Dec;104(6):3476–93.

25. Pennartz CMA, Uylings HBM, Barnes CA, McNaughton BL. Memory reactivation and consolidation during sleep: from cellular mechanisms to human performance. Prog Brain Res. 2002;138:143–66.

26. Horovitz SG, Braun AR, Carr WS, Picchioni D, Balkin TJ, Fukunaga M, et al. Decoupling of the brain’s default mode network during deep sleep. Proc Natl Acad Sci U S A. 2009 Jul 7;106(27):11376–81.

27. Girardeau G, Benchenane K, Wiener SI, Buzsáki G, Zugaro MB. Selective suppression of hippocampal ripples impairs spatial memory. Nat Neurosci. 2009 Oct;12(10):1222–3.

28. Maingret N, Girardeau G, Todorova R, Goutierre M, Zugaro M. Hippocampo-cortical coupling mediates memory consolidation during sleep. Nat Neurosci. 2016 Jul;19(7):959–64.

29. Barthó P, Hirase H, Monconduit L, Zugaro M, Harris KD, Buzsáki G. Characterization of neocortical principal cells and interneurons by network interactions and extracellular features. J Neurophysiol. 2004 Jul;92(1):600–8.

30. Massimini M, Huber R, Ferrarelli F, Hill S, Tononi G. The sleep slow oscillation as a traveling wave. J Neurosci. 2004 Aug 4;24(31):6862–70.

31. Mölle M, Yeshenko O, Marshall L, Sara SJ, Born J. Hippocampal sharp wave-ripples linked to slow oscillations in rat slow-wave sleep. J Neurophysiol. 2006 Jul;96(1):62–70.

32. Vyazovskiy VV, Riedner BA, Cirelli C, Tononi G. Sleep homeostasis and cortical synchronization: II. A local field potential study of sleep slow waves in the rat. Sleep. 2007 Dec;30(12):1631–42.

33. Skaggs WE, McNaughton BL. Replay of neuronal firing sequences in rat hippocampus during sleep following spatial experience. Science. 1996 Mar 29;271(5257):1870–3.

34. Kim S, Putrino D, Ghosh S, Brown EN. A Granger causality measure for point process models of ensemble neural spiking activity. PLoS Comput Biol. 2011 Mar;7(3):e1001110.

35. Vinck M, Huurdeman L, Bosman CA, Fries P, Battaglia FP, Pennartz CMA, et al. How to detect the Granger-causal flow direction in the presence of additive noise? NeuroImage. 2015 Mar;108:301–18.

36. Quian Quiroga R, Panzeri S. Extracting information from neuronal populations: information theory and decoding approaches. Nat Rev Neurosci. 2009 Mar;10(3):173–85.

37. Schreiber. Measuring information transfer. Phys Rev Lett. 2000 Jul 10;85(2):461–4.

38. Barnett L, Barrett AB, Seth AK. Granger causality and transfer entropy are equivalent for Gaussian variables. Phys Rev Lett. 2009 Dec 4;103(23):238701.

39. Harris KD, Thiele A. Cortical state and attention. Nat Rev Neurosci. 2011 Sep;12(9):509–23.

40. Sirota A, Csicsvari J, Buhl D, Buzsáki G. Communication between neocortex and hippocampus during sleep in rodents. Proc Natl Acad Sci U S A. 2003 Feb 18;100(4):2065–9.

41. Rasch B, Born J. About sleep’s role in memory. Physiol Rev. 2013 Apr;93(2):681–766.

42. Buzsáki G. Two-stage model of memory trace formation: a role for “noisy” brain states. Neuroscience. 1989;31(3):551–70.

43. Koch C, Massimini M, Boly M, Tononi G. Neural correlates of consciousness: progress and problems. Nat Rev Neurosci. 2016 Apr 20;17(5):307–21.

44. Pennartz CMA. The Brain’s Representational Power [Internet]. MIT Press; 2015 [cited 2017 Aug 29]. Available from: https://mitpress.mit.edu/books/brains-representational-power

45. Storm JF, Boly M, Casali AG, Massimini M, Olcese U, Pennartz CMA, et al. Consciousness Regained: Disentangling Mechanisms, Brain Systems, and Behavioral Responses. J Neurosci. 2017 Nov 8;37(45):10882–93.

46. Hoffman KL, McNaughton BL. Coordinated reactivation of distributed memory traces in primate neocortex. Science. 2002 Sep 20;297(5589):2070–3.

47. Rothschild G, Eban E, Frank LM. A cortical-hippocampal-cortical loop of information processing during memory consolidation. Nat Neurosci. 2017 Feb;20(2):251–9.

48. Qin YL, McNaughton BL, Skaggs WE, Barnes CA. Memory reprocessing in corticocortical and hippocampocortical neuronal ensembles. Philos Trans R Soc Lond B Biol Sci. 1997 Oct 29;352(1360):1525–33.

49. Pennartz CMA, Lee E, Verheul J, Lipa P, Barnes CA, McNaughton BL. The ventral striatum in off-line processing: ensemble reactivation during sleep and modulation by hippocampal ripples. J Neurosci. 2004 Jul 21;24(29):6446–56.

50. Sato TK, Nauhaus I, Carandini M. Traveling waves in visual cortex. Neuron. 2012 Jul 26;75(2):218–29.

51. Compte A, Sanchez-Vives MV, McCormick DA, Wang X-J. Cellular and network mechanisms of slow oscillatory activity (<1 Hz) and wave propagations in a cortical network model. J Neurophysiol. 2003 May;89(5):2707–25.

52. Olcese U, Faraguna U. Slow cortical rhythms: from single-neuron electrophysiology to whole-brain imaging in vivo. Arch Ital Biol. 2015 Sep;153(2–3):87–98.

53. Piarulli A, Menicucci D, Gemignani A, Olcese U, d’Ascanio P, Pingitore A, et al. Likeness-based detection of sleep slow oscillations in normal and altered sleep conditions: application on low-density EEG recordings. IEEE Trans Biomed Eng. 2010 Feb;57(2):363–72.

54. Dupret D, O’Neill J, Pleydell-Bouverie B, Csicsvari J. The reorganization and reactivation of hippocampal maps predict spatial memory performance. Nat Neurosci. 2010 Aug;13(8):995–1002.

55. Schönauer M, Geisler T, Gais S. Strengthening procedural memories by reactivation in sleep. J Cogn Neurosci. 2014 Jan;26(1):143–53.

56. Girardeau G, Zugaro M. Hippocampal ripples and memory consolidation. Curr Opin Neurobiol. 2011 Jun;21(3):452–9.

57. Jadhav SP, Kemere C, German PW, Frank LM. Awake hippocampal sharp-wave ripples support spatial memory. Science. 2012 Jun 15;336(6087):1454–8.

58. Diba K, Buzsáki G. Forward and reverse hippocampal place-cell sequences during ripples. Nat Neurosci. 2007 Oct;10(10):1241–2.

59. Pfeiffer BE, Foster DJ. Hippocampal place-cell sequences depict future paths to remembered goals. Nature. 2013 May 2;497(7447):74–9.

60. Roumis DK, Frank LM. Hippocampal sharp-wave ripples in waking and sleeping states. Curr Opin Neurobiol. 2015 Dec;35:6–12.

61. Casali AG, Gosseries O, Rosanova M, Boly M, Sarasso S, Casali KR, et al. A theoretically based index of consciousness independent of sensory processing and behavior. Sci Transl Med. 2013 Aug 14;5(198):198ra105.

62. Vinck M, Bos JJ, Van Mourik-Donga LA, Oplaat KT, Klein GA, Jackson JC, et al. Cell-Type and State-Dependent Synchronization among Rodent Somatosensory, Visual, Perirhinal Cortex, and Hippocampus CA1. Front Syst Neurosci. 2015;9:187.

63. Bos JJ, Vinck M, van Mourik-Donga LA, Jackson JC, Witter MP, Pennartz CMA. Perirhinal firing patterns are sustained across large spatial segments of the task environment. Nat Commun. 2017 May 26;8:15602.

64. Wang Q, Chen F, Xu W, Yang M-H. Object Tracking via Partial Least Squares Analysis. IEEE Trans Image Process. 2012 Oct;21(10):4454–65.

65. Iurilli G, Olcese U, Medini P. Preserved excitatory-inhibitory balance of cortical synaptic inputs following deprived eye stimulation after a saturating period of monocular deprivation in rats. PloS One. 2013;8(12):e82044.

66. Olcese U, Iurilli G, Medini P. Cellular and synaptic architecture of multisensory integration in the mouse neocortex. Neuron. 2013 Aug 7;79(3):579–93.

67. Saleem AB, Chadderton P, Apergis-Schoute J, Harris KD, Schultz SR. Methods for predicting cortical UP and DOWN states from the phase of deep layer local field potentials. J Comput Neurosci. 2010 Aug;29(1–2):49–62.

68. Vicente R, Wibral M, Lindner M, Pipa G. Transfer entropy--a model-free measure of effective connectivity for the neurosciences. J Comput Neurosci. 2011 Feb;30(1):45–67.

69. Magri C, Whittingstall K, Singh V, Logothetis NK, Panzeri S. A toolbox for the fast information analysis of multiple-site LFP, EEG and spike train recordings. BMC Neurosci. 2009;10:81.

